## Supplemental for "Endophilin A2 regulates B cell protein trafficking and humoral responses"

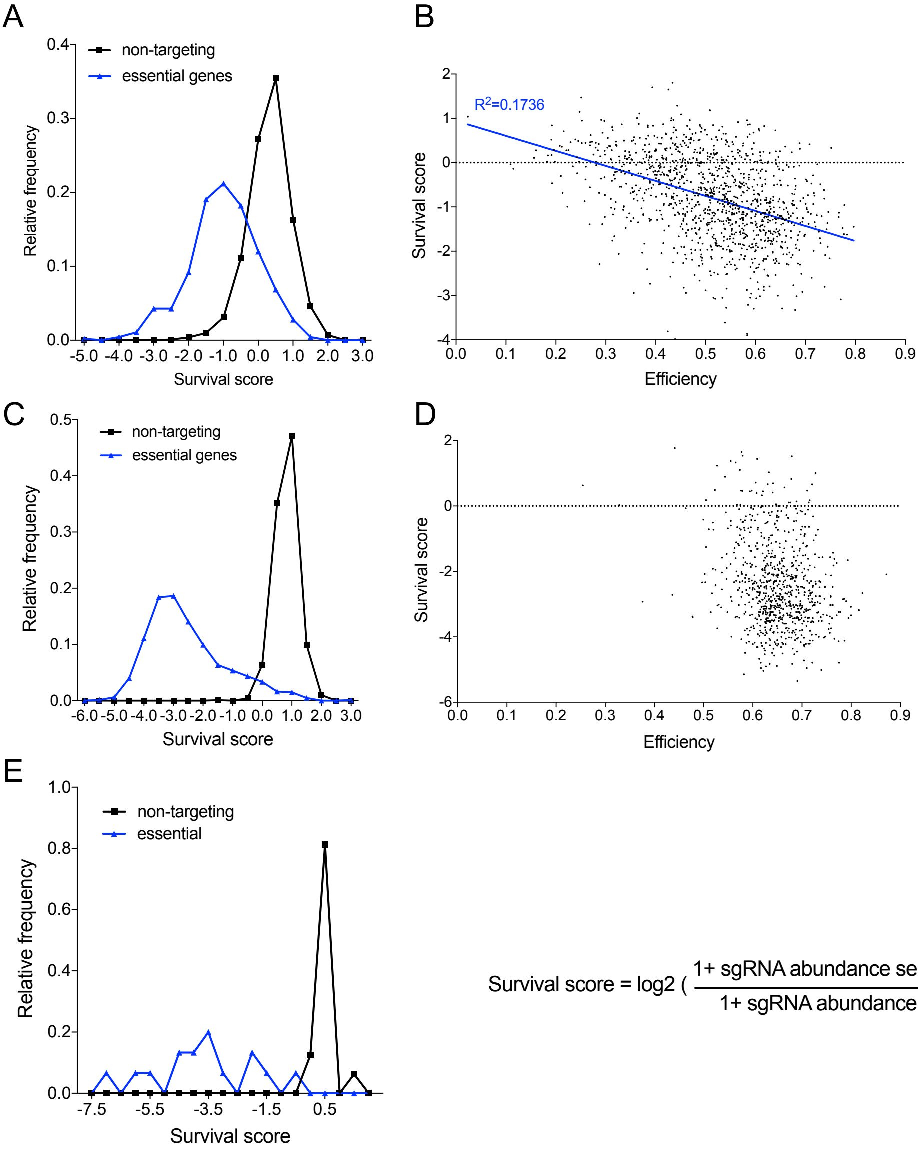


$$\text{Survival score} = \log 2\left(\frac{1+\text{ sgRNA abundance selected population}}{1+\text{ sgRNA abundance initial infection}}\right)$$

Figure S1. Analysis of essential gene targeting in large-scale pooled CRISPR screens. (A) GeCKO library essential gene survival scores. The plot shows mean CRISPR survival scores of sgRNAs targeting a set of top 200 essential genes (Wang et al. 2015). (B) Correlation of the CRISPR survival score and sgRNA on-target efficiency for a set of sgRNAs targeting the essential genes from the GeCKO library. On-target efficiency was calculated using “Rule Set 2” algorithm (Doench et al. 2016). (C) Brunello library essential gene survival scores. (D) Correlation of survival score and sgRNA on-target efficiency in Brunello library. (E) Custom minilibrary essential gene survival scores.

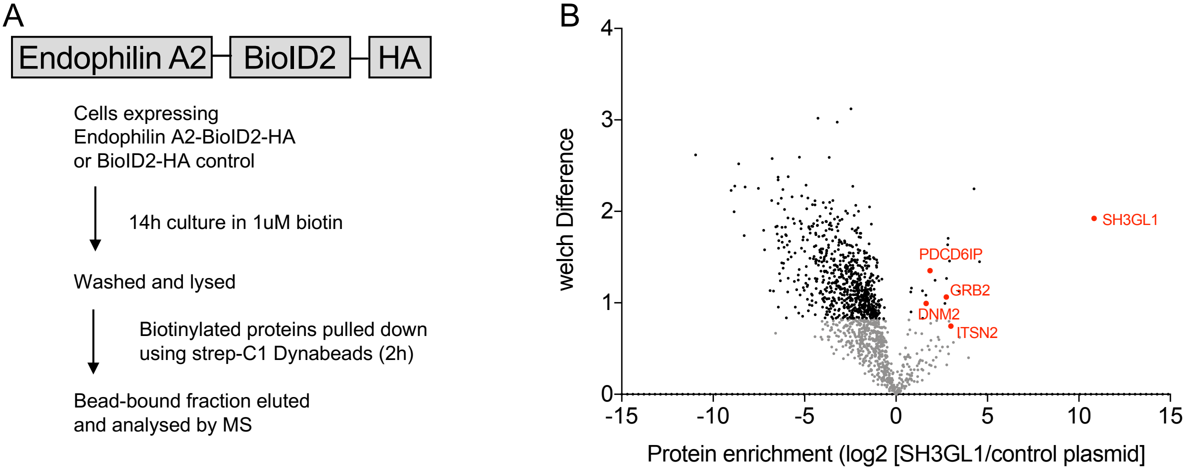


Figure S2. BioID2 reveals potential endophilin A2 binding partners. (A) BioID2-tagged endophilin A2 construct and assay workflow. (B) Proteins significantly enriched in biotinylated fraction of endophilin A2-BioID2 samples compared to empty plasmid control (positive protein enrichment scores). Negative enrichment score represents gene products enriched in BioID2 control sample, reflecting the promiscuous nature of the cytoplasmic biotinylase. Data from 2 independent experiments, see also Supplemental table 4.

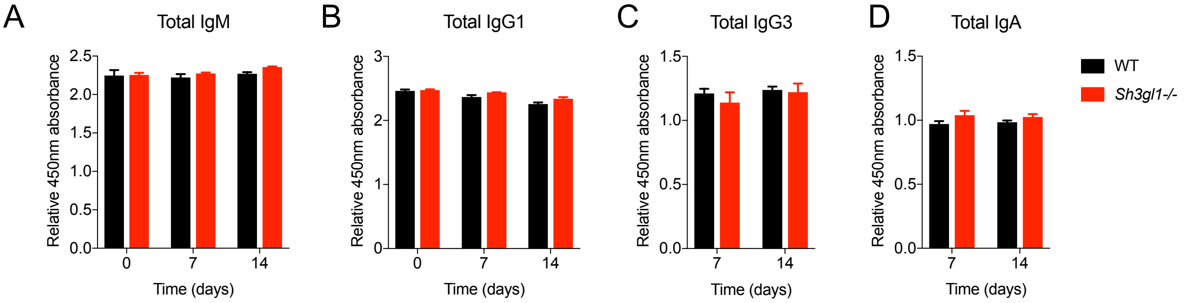


Figure S3. Total antibody titers are unaffected in *Sh3gl1*$^{-/-}$ mice. (A-D) Total antibody levels measured by clonotyping ELISA using capture antibody. One representative experiment showing measurements from 4 mice. (A) IgM (B) IgG1 (C) IgG3 (D) IgA.

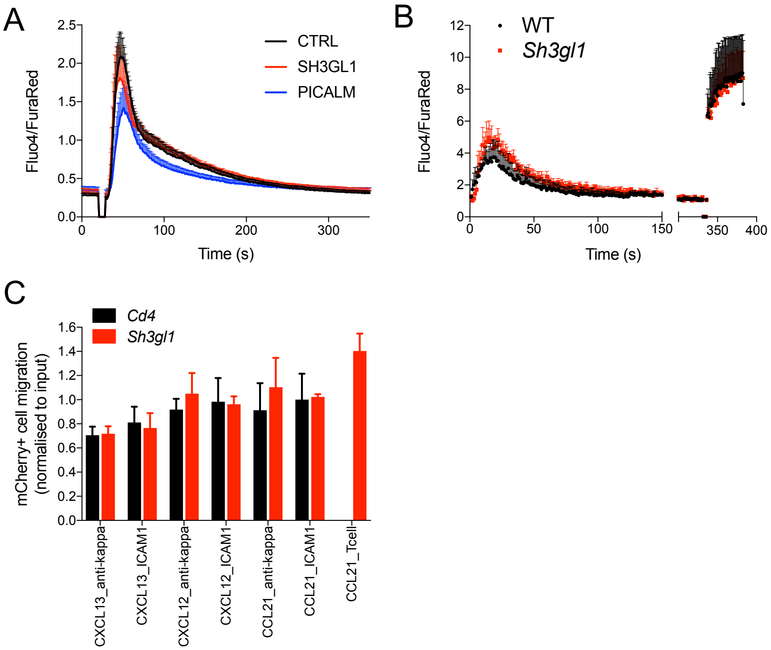


Figure S4. Normal antigen-induced calcium signaling and chemokine-induced migration in the absence of endophilin A2. (A) CRISPR-targeted Ramos, activated with 2 µg/ml of anti-human IgM $F(ab')_2$. Data show mean and SEM from 4 independent infections. (B) Primary B cells from *Sh3gl1*$^{-/-}$ mice or WT littermates, activated with 2 µg/ml anti-mouse Igκ $F(ab')_2$, for 380 seconds, followed by addition of ionomycin (5 µg/ml) at 300 s. N = 3 mice. (C) Chemokine-induced migration through ICAM-1 or anti-Igκ coated transwells. mCherry$^+$ percentage is normalized to input population.

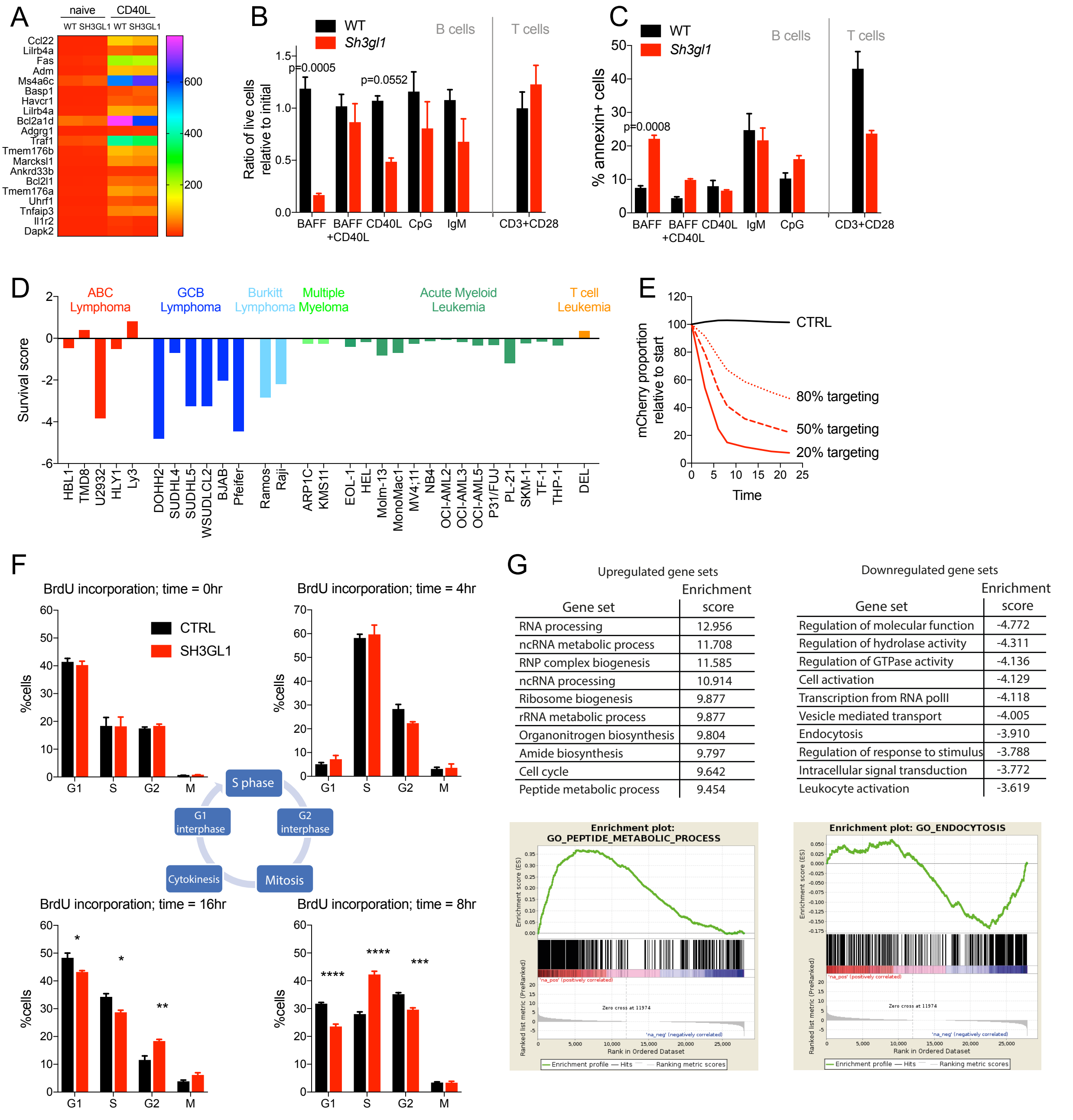


Upregulated gene sets

| Gene set | Enrichment score |
|---|---|
| RNA processing | 12.956 |
| ncRNA metabolic process | 11.708 |
| RNP complex biogenesis | 11.585 |
| ncRNA processing | 10.914 |
| Ribosome biogenesis | 9.877 |
| rRNA metabolic process | 9.877 |
| Organonitrogen biosynthesis | 9.804 |
| Amide biosynthesis | 9.797 |
| Cell cycle | 9.642 |
| Peptide metabolic process | 9.454 |

Downregulated gene sets

| Gene set | Enrichment score |
|---|---|
| Regulation of molecular function | -4.772 |
| Regulation of hydrolase activity | -4.311 |
| Regulation of GTPase activity | -4.136 |
| Cell activation | -4.129 |
| Transcription from RNA polII | -4.118 |
| Vesicle mediated transport | -4.005 |
| Endocytosis | -3.910 |
| Regulation of response to stimulus | -3.788 |
| Intracellular signal transduction | -3.772 |
| Leukocyte activation | -3.619 |

Figure S5. Loss of endophilin A2 affects cell expansion in primary and Ramos B cells. (A) Heat map of TPM values from RNAseq comparing follicular B cells from WT and *Sh3gl1*$^{-/-}$ littermates. Shown are the top 20 genes upregulated in WT B cells 24 hours after CD40L activation. (B) Numbers of viable cells in 3-day culture of follicular B cells in specified cytokines. N = 3 mice. (C) AnnexinV stain of B cell cultures in B. (B, C) Data show mean and SEM; P values are calculated using two-way ANOVA. (D) Endophilin A2 survival scores of Ramos cells using the Brunello library, together with CRISPR scores from published genome-wide screens in hematopoietic cell lines (Phelan et al. 2018; Wang et al. 2017; Wang et al. 2015). Scores are normalized using the mean value of the top 200 essential genes in each screen. (E) Decreased percentage of *SH3GL1*-targeted Ramos population in experiments with indicated initial percentage of targeted cells. Data show one representative experiment out of 3. (F) BrdU pulse chase assay in Ramos cells. Clockwise from top left: equal BrdU incorporation at time = 0h; slower progression of BrdU-labelled cells into G2 phase in *SH3GL1*- compared to CTRL- targeted cells at time = 4h; slower progression into G1 of following cycle while greater proportion of *SH3GL1* cells remain in initial S phase at time = 8h; slower progression between G2 and M phase in *SH3GL1*-targeted cells resulting in significantly greater numbers in G2 and less in G1 or S phase of subsequent cycle. (G) GSEA of RNAseq analysis comparing WT and *Sh3gl1*$^{-/-}$ sorted follicular B cells. 3 mice of each genotype were used for RNAseq and top 10 significantly enriched gene sets up- or down-regulated in *Sh3gl1*$^{-/-}$ B cells are shown.

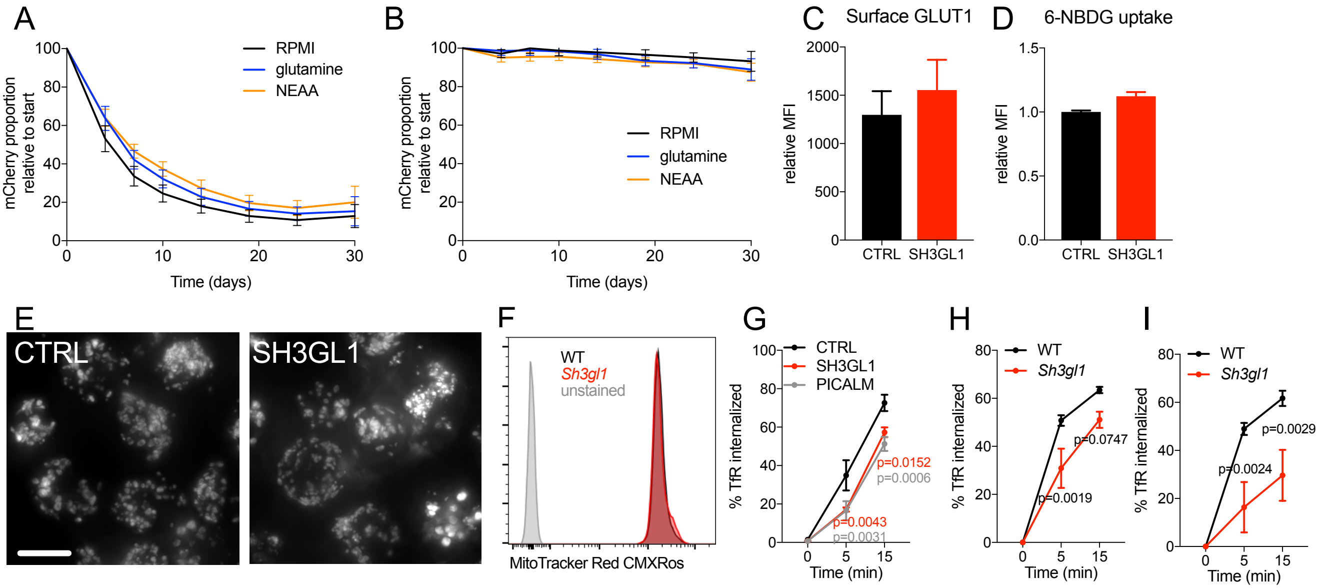


Figure S6. Growth defect upon *SH3GL1* deletion is not rescued by common cell culture supplements and does not correlate with altered mitochondrial mass. (A) mCherry percentage in *SH3GL1*-targeted Ramos cells over time, supplemented with 5x normal concentration of L-Glutamine or NEAA. (B) mCherry percentage in CTRL-targeted Ramos cells over time, supplemented with 5x normal concentration of L-Glutamine or NEAA. (C) Surface GLUT1 stain in CRISPR-targeted Ramos cells, N = 8 across 3 experiments. (D) Accumulation of 6-NBDG (fluorescent glucose analog) after 10min internalization in Ramos cells; N = 4. (E) Anti-TUFM immunofluorescence showing total mitochondrial mass and organization in CRISPR-targeted Ramos cells. (F) MitoTracker Red CMXRos stain for active mitochondria in primary B cells from WT and *Sh3gl1*$^{-/-}$ B cells. (G) TFRC internalization assay in Ramos cells using biotinylated anti-CD71. Data show mean ± SEM from 4 independent CRISPR infections in 2 experiments. P, significance calculated using two-way ANOVA and indicated in corresponding color. (H, I) Mouse transferrin internalization in B cells from SRBC-immunized WT and *Sh3gl1*$^{-/-}$ littermates. Data shows mean and SEM from 7 mice per genotype in 2 experiments. P, statistical significance analyzed using two-way ANOVA. (H) Naïve B cells. (I) GC B cells.
